# Cues for predictive eye movements in naturalistic scenes

**DOI:** 10.1101/2023.04.21.537766

**Authors:** Alexander Goettker, Nils Borgerding, Linus Leeske, Karl R. Gegenfurtner

**Affiliations:** Justus Liebig Universität Giessen, Germany; Center for Mind, Brain and Behavior, University of Marburg and Justus Liebig University

## Abstract

We previously compared following of the same trajectories with eye movements, but either as an isolated target or embedded in a naturalistic scene, in this case the movement of a puck in an ice hockey game. We observed that the oculomotor system was able to leverage the contextual cues available in the naturalistic scene to produce predictive eye movements. In this study we wanted to assess which factors are critical for achieving this predictive advantage by manipulating four factors: the expertise of the viewers, the amount of available peripheral information, and contextual and kinematic cues. The more peripheral information became available (by manipulating the area of the video that was visible), the better the predictions of all observers. However, expert ice hockey fans were consistently more accurate and better at predicting than novices and also benefitted more from additional peripheral information. Artificial contextual cues about the position of the players did not lead to a predictive advantage, whereas impairing the causal structure of kinematic cues by playing the video in reverse led to a severe impairment. When the videos were flipped vertically to introduce more difficult kinematic cues, predictive behavior was comparable to when observers watching the original videos. Together, these results demonstrate that when contextual information is available in naturalistic scenes, the oculomotor system is successfully integrating them, and is not only relying on low-level information about the target trajectory. Critical factors for successful prediction seem to be the amount of available information, experience with the stimuli and the availability of intact kinematic cues for player movements.

## Introduction

Over the last decades eye movements became widely used as a window into the brain, mind, and cognitive processes (König et al., 2016; Shaikh & Zee, 2018). The success of eye movement research is related to results that demonstrate a direct link between basic oculomotor paradigms, such as the antisaccade task (Hallett, 1978; for reviews see Hutton & Ettinger, 2006; Munoz & Everling, 2004) or saccadic adaptation (McLaughlin, 1967; for a review see Pélisson et al., 2010), and different cognitive metrics or learning mechanisms. In addition, the continuous nature of eye movement responses even allows one to map the dynamics of neuronal representations (Pack & Born, 2001). However, one thing that most of these influential paradigms have in common is the use of highly-controlled, but very simplistic stimuli (e.g., a simple target disk). Classically, oculomotor control is assumed to be mainly driven by early stages of visual processing (Lisberger, 2010, 2015) manipulated in such paradigms. However, multiple studies revealed that higher-level factors such as the task (Chen et al., 2021; Rothkopf et al., 2016; Tatler et al., 2011; Yarbus, 1967) or expectations (Diaz et al., 2013; Jörges & López-Moliner, 2019; Kowler et al., 2019; Vo & Henderson, 2009) also have a strong impact on how we move our eyes. Therefore, a critical question remains unanswered: Can effects and mechanisms identified with these simple stimuli be generalized to eye movement behavior with more naturalistic stimuli and tasks?

The most influential early work on eye movement behavior in naturalistic settings was performed when using mobile eye trackers to record eye movement during everyday tasks (Hayhoe & Ballard, 2005; Land & Hayhoe, 2001). In these studies, eye movements were recorded while observers prepared a tea or making a peanut-butter jelly sandwich. One of the key results of these studies was that observes mostly fixated on objects they interacted with, but already moved their gaze predictively to the next relevant object before they completed the preceding action. This suggests that an important goal of the oculomotor system is to bring gaze to relevant locations at the right time to gather information. However, this is not a trivial task, since there are substantial internal processing delays (Nowak & Bullier, 1997; Schlag & Schlag-Rey, 2002). For simple stimuli, we know that despite these processing delays it is possible to make accurate saccades to moving targets (Fleuriet et al., 2011; Goettker et al., 2019; Ron et al., 1989; Schreiber et al., 2006) or to keep track of a predictably moving target without lag (Terry Bahill & McDonald, 1983). For naturalistic stimuli, close to zero delay tracking has been observed when watching videos of outdoor scenes (Vig et al., 2011). However, under more naturalistic contexts, predictive eye movements seem to occur more frequently, which can either be driven by knowledge about the physics of the world (Diaz et al., 2013), monitoring and planning a task (Sullivan et al., 2021), or the semantic information provided by the scene (Henderson, 2017). However, until recently a direct comparison of oculomotor behavior for simple and naturalistic stimuli was missing.

In our recent study (Goettker et al., 2021) we tried to fill this gap. We made use of a hand-labelled data set of ice hockey videos (Pidaparthy & Elder, 2019), which allowed us to present the movement of the puck (represented by a disk) in front of a gray background. We compared this condition with the original videos. For both conditions the task of the observers was to follow the puck movements with their eyes and due to the identical trajectories, the low-level motion characteristics were comparable. When we computed the delay that observers had when tracking the motion trajectory, we observed a striking difference: With only the target movement in front of a gray background, the estimated delay was around 180 ms, which is expected with purely reactive behavior. In contrast, when seeing the original videos, even our non-expert observers had an average delay close to zero. This suggests that observers were able to make use of complex naturalistic input and integrated it to significantly reduce their tracking delay. However, since the conditions in our previous study were the two extreme cases of either no additional information present (when just seeing the disk) or all additional information available (original video), it is unclear which cues and information observers used to achieve this benefit in tracking performance. The goal of the current study is to answer that question by manipulating four factors: (1) the level of expertise of the observers, (2) the amount of peripheral information available, and the availability and reliability of (3) contextual and (4) kinematic cues.

In our original study, we had studied non-expert observers and even they reached almost zero-delay tracking when watching hockey videos. Therefore, we wanted to investigate how experts perform in our task, since it is known that across many different aspects (Brams et al., 2019), and especially in sports (Memmert, 2009; Vickers, 2009), expertise leads to differences in behavior and performance. We combined the difference in expertise with the investigation of the role of the amount of peripheral information available. We achieved this by using manipulated versions of the videos, where only a certain area around the puck position was visible. This is of interest due to two reasons: First, can experts make better use of additional information by better integration them for oculomotor control (Brams et al., 2019; Casanova et al., 2009). Second, additional information might also be related to more crowding and limits in peripheral processing (Rosenholtz, 2016; Rosenholtz et al., 2007; Whitney & Levi, 2011), therefore additional peripheral information could also lead to no difference or even worse performance. For the other factor of interest, we used a second experiment in which we manipulated information along two axis that are known to be helpful in anticipation: context and kinematic cues (Loffing & Cañal-Bruland, 2017). For contextual cues, we added information about the player positions (comparable to the classical: Heider & Simmel, 1944), but did not show the kinematic cues of player movements. To manipulate kinematic cues, we kept the video intact, but played it in reverse or flipped it vertically to reduce their reliability (Pavlova et al., 2002; Pavlova & Sokolov, 2000). Together, the combination of these different manipulations should give us a good starting point to understand which information observers used to gain the predictive advantage in the video condition.

## Experiment 1 - The role of expertise and peripheral information

### Methods

#### Observer

26 observers took part in the experiment. Half of the participants were expert viewers, defined by reporting a good understanding of tactical knowledge in ice hockey games and the self report of regularly watching ice hockey games (on average 1.88 +/-1.29 games per week), and were fans of the sport since for at least a year. In contrast, the novice group never watched an ice hockey game before. With respect to other characteristics such as gender (in both groups there were 4 persons who identified as female, 9 who identified as male) or age (M = 23.46 +/-1.71 for the experts, M = 23.31 +/-1.75) the groups were matched. All participants were naïve with respect to the purpose of the study and gave informed consent at the beginning of the experiment. The experiment was in line with the Declaration of Helsinki and was approved by the local ethics committee. Participants were paid for their effort with 8 euros/hour.

#### Setup

Participants sat at a table in a dimly illuminated room with their head positioned on a chin rest. In this position, their eyes were roughly at the height of the center of a monitor (3840 × 2160 pixel, Phillips, Amsterdam, Netherlands) at a distance of 70 cm. In this setup the monitor spanned approximately 49 × 26 deg of visual angle. The experiment was programmed and controlled with Matlab (MathWorks, Natick MA) using Psychtoolbox(Kleiner et al., 2007). Gaze was recorded from one eye with a desk-mounted eye tracker (EyeLink 1000 Plus, SR Research, Kanata, ON, Canada) at a sampling frequency of 1000 Hz. Before each block a nine-point calibration was used, and an additional drift check was performed at the start of each trial.

#### Stimuli & Conditions

We selected 12 of the originally 18 videos (each lasting for 10s) from our previous paper (Goettker et al., 2021) for the follow up experiments. The selection was based on taking videos that contained the labelled passes and showed the most reliable effects in the original study. Please note here, that all videos were filmed with a static camera, therefore no artifacts due to camera movements were present and observers had to keep track of the puck just by moving their eyes. In addition to the expert/novice group, we created four different conditions for each of the videos which systematically varied the amount of peripheral information that was available. Two conditions were identical to our previous experiment: The *disk condition* was just a small black disk (diameter = 0.3 deg) moving along the labelled puck position (see original paper for more details) in front of uniform gray background and therefore no peripheral information was available. In the *video condition* the original, unaltered hockey clip was shown and therefore all available peripheral information was presented. We created two intermediate conditions, where we only showed a cutout of the video around the labelled puck position. We created the cutouts by using a 2D Gaussian centered on the puck position, which was by multiplied with the original images and added a uniform gray background image multiplied by 1-Gaussian weights. We used a standard deviation of 3 deg as *small context condition* and a standard deviation of 8 deg as *large context condition*.

Importantly, participants were explicitly told that their task is to follow the target (a disk in the disk condition or the puck in the rest of the conditions). Each video was shown as a single trial. Each trial started with a central fixation cross, which was used for a drift check. Observers started the trial by looking at the cross and pressing the space bar at their own pace. Then the video played for 10 s. After the video ended a gray screen appeared, and a message appeared that indicated that the next trial is loading. The message stayed on the screen for roughly 5 s, in which the next video was loaded into Psychtoolbox to prepare for the next trial.

All observers completed all four conditions in a random order. The order was matched between the expert and novice group such that each recruited expert saw the conditions in the same order as one of the recruited novices. In each condition observers completed three blocks and in each block all videos were shown in a random order. Each participant completed 144 trials (4 conditions x 3 blocks x 12 videos). One block took roughly 5-7 minutes and participants completed all 12 blocks in one 1.5-hour session. Across all participants this led to a total of 3744 trials (2 groups x 13 observers x 144 trials).

#### Data Analysis

Raw gaze positions were saved during each trial and analyzed offline with custom analysis scripts in MATLAB (MathWorks, Natick MA). Blinks in the data were linearly interpolated. Gaze position data was filtered with a second-order Butterworth filter with a cutoff frequency of 30 Hz. Afterwards, eye velocity was computed based on the difference in horizontal and vertical eye position in consecutive samples, which then was converted into the absolute 2D eye velocity in deg/s by taking the square root of the squared differences and multiplying it by the sampling rate. Since the target position was only available at the frame rate of the video (30 Hz), it was upsampled via linear interpolation to match the sampling rate of eye position. Saccades were detected based on the EyeLink criteria (velocity > 30 deg/s & acceleration > 4000 deg/s^2^). Following that, for each trial we extracted different metrics related to different aspects of tracking performance.

For overall performance we computed the tracking error as the median Euclidean distance between gaze and puck position. We removed the first 500 ms from the analysis, since in the beginning of each trial participants needed to fixate in the center of the screen and initially always had to search for the target. We also computed the number of saccades, and the proportion of the video spend in pursuit (defined as eye velocity > 3 deg/s for at least 100 ms with no saccade) as general measures of tracking performance.

To estimate the delay between eye and target movement we used a cross-correlation approach. We used a sliding window with a width of 500 ms. For each window, we computed the correlation between eye and target position for varying delays (−200 to +400 ms in steps of 10 ms) separately for horizontal and vertical eye positions. Then the window was shifted by 250 ms and the procedure repeated until the end of the video was reached. At the end of the video the median correlation across the different sliding windows for each delay was computed. To account for auto-correlation within the data, we estimated the correlation within each clip by performing 1000 samples for 500 ms windows for eye and puck at random time points (for example the eye vector after 1s and the target vector after 8s) again for horizontal and vertical eye position. We then subtracted those from the median. Finally, we took the average of the correlations obtained for horizontal and vertical movements, to obtain one correlation value for each of the different delays between gaze and puck/disk movement for a given trial. This approach allowed us to quantify the delay between eye and target movement independent of the specific oculomotor response.

We also estimated the delay separately for saccadic and pursuit eye movements. For pursuit eye movements we took a very similar approach, however instead of using a sliding window across the whole video, we performed the cross-correlation approach for each pursuit segment found in the data. Pursuit segments were identified as time windows of at least 100 ms, where the eye velocity was above 3 deg/s and no saccade occurred. We did not estimate a pursuit delay when the overall proportion of pursuit for a respective observer and condition was below 0.2, since then too few pursuit estimates were available. To estimate the delay for each saccade, we projected the saccade endpoint on a straight line connecting the target position 100 ms before saccade onset and saccade end. Then, we computed the distance between the projected endpoint and the target position 100 ms before saccade onset and took the ratio between this value and the actual distance the target moved during that time. Therefore, a ratio above 1 was related to an overshoot of the target (predictive to the future), whereas a ratio below 1 was related to a undershoot (lag behind the target). To compare it with the estimated delay in ms for pursuit eye movements, we turned that ratio into a ms estimate by multiplying it with the duration between 100 ms before saccade onset and saccade offset. In a last step we normalized this value by subtracting the respective time again, so that a ratio of 1 led to an estimate of 0 ms delay. Across all conditions and subjects, eye movement data consisted of 9% saccades and 32% pursuit, with the remainder being considered as fixations, including small drift movements.

As a special example to visualize predictions, we analyzed 11 hand-labelled passed between players in more detail. As a measurement of prediction, we estimated the time of gaze arrival at the targeted player. We searched for the first instance around the time of the pass when gaze was closer than 3 deg to the final puck position. To control for the residual time that the puck still needed towards the target location, we also measured the time the puck arrived within 3 deg of the final location and subtracted this value from the estimated time of gaze arrival. To assess the accuracy of the prediction, we computed the error after the pass. We calculated the median gaze position between 200 and 300 ms after the pass offset and projected it onto the pass trajectory. The comparison of the actual endpoint and the projected endpoint gave us the error and the direction of the error was reflected in an overshoot or undershoot of the target position after the end of the pass.

#### Exclusion criteria and statistical analysis

Trials were excluded from the analysis based on two criteria: (1) if during the trial data was missing for more than 500 ms in a row and (2) the computed average error was larger than 10 deg. Based on these criteria 3575 of 3744 trials (96 %) were included in the analysis.

For each measure the average across all 12 videos across all 3 blocks was computed for each subject, so that there was one value per subject per condition. The obtained values were based on averages across values from each video (e.g. position error), each pass (e.g. time to arrive at pass target), or all saccades and pursuit segments that were present across all videos from one condition (estimate of saccade & pursuit delay). The averages were computed and then statistical analysis was performed in JASP. Participants were grouped by their expertise and repeated-measure ANOVAs with the factor Peripheral Information (Disk, Small, Large, Video) and the inter-subject factor Expertise (Novice, Expert) was computed. If the sphericity assumption was violated, Greenhouse-Geisser correction was used to correct the degrees of freedom, and the corrected values are reported. Post-hoc t-tests were used to assess differences between the conditions and to correct for multiple testing the adjusted p-value according to Holm was reported. The default option of pooled error terms for RM factors was kept active.

### Results

The goal of experiment 1 was to investigate the influence of expertise as well as the amount of available peripheral information. To quantify oculomotor behavior, we used different metrics; we will start with reporting measurements quantifying changes in overall eye movement behavior, followed by computations of the delay between eye and target movements. In the last step, we will take a closer look at eye movement behavior in pass situations, as they are a prime example to visualize predictive behavior.

#### Description of overall behavior

To quantify overall tracking behavior, we looked at three metrics: position error, number of saccades and proportion of pursuit eye movements (see Methods for details). We assessed the influence of our manipulated variables with a repeated-measurement ANOVA with the factor peripheral information (Disk, Small, Large, Video) and the between-subject factor Expertise (Novice vs Expert). For position error we observed a significant main effect of the available peripheral information (F(1.583, 37.995) = 38.908, p < .001) as well a main effect of expertise (F(1,24) = 5.405, p = .029), but no significant interaction (F(1.583, 37.995) = 0.520, p = .557). Experts had significantly lower error across all conditions (see Figure 2 A), but the position error significantly increased with increasing peripheral information (all comparisons p <.001, except disk vs. small p = .162) with the highest position error when tracking the puck in the full video. This suggests a slight increase in position error when more complex visual information was available, which might have made it more difficult to find the puck or could suggest a different tracking strategy.

**Figure 1.**
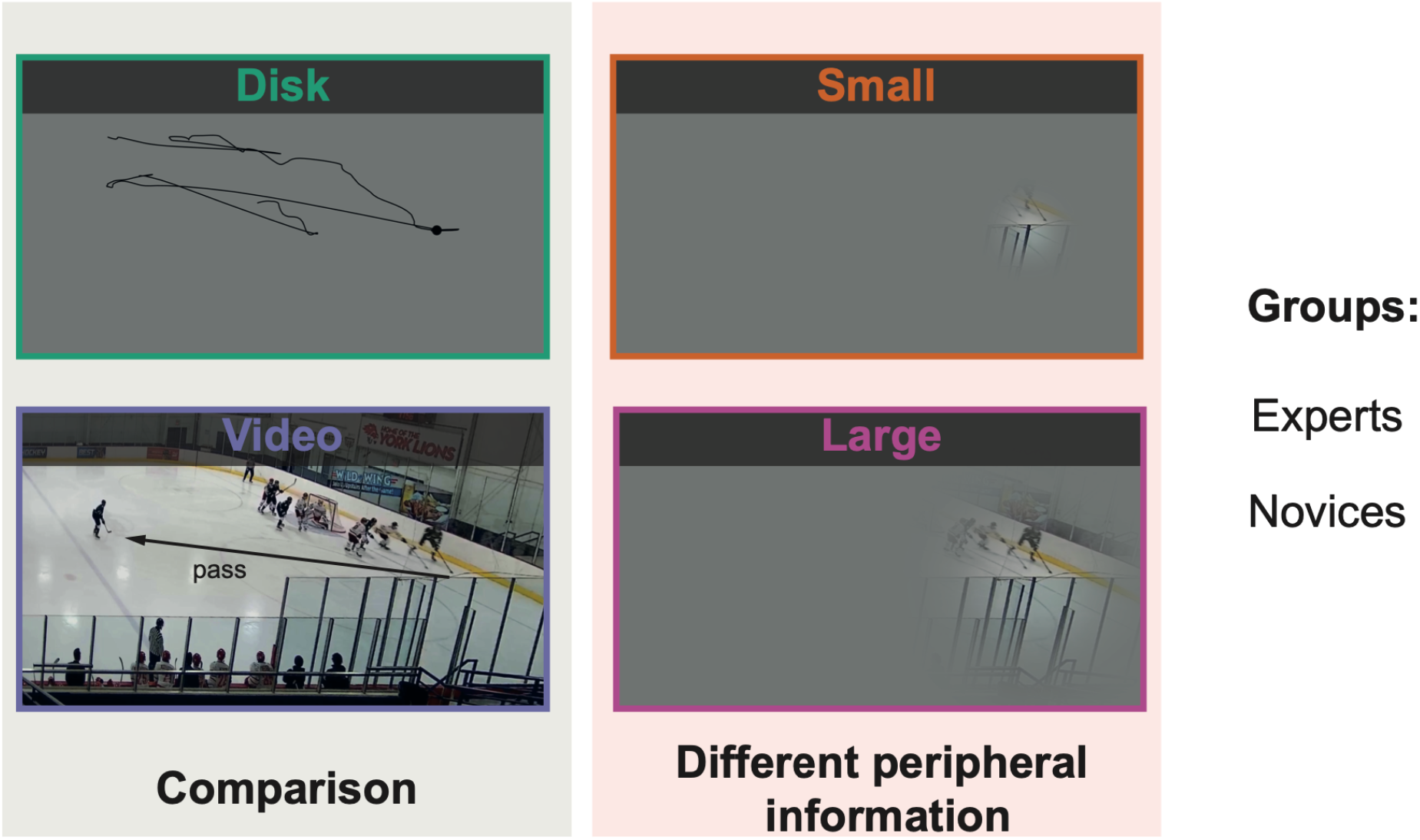
Conditions of Experiment 1. Depiction of the four different conditions. Disk and video conditions are the conditions previously tested in Goettker et al. 2021. In the disk condition the whole trajectory of the puck is illustrated as an example, please note that this was never visible to the observer. For the video condition, a pass between two players is illustrated, since these situations will be one focus of the analysis. The small and large context condition manipulate the available peripheral information by only showing certain cutouts around the pass position. All four conditions were performed by each observer, but observers were either experts or novices with respect to ice hockey.

**Figure 2.**
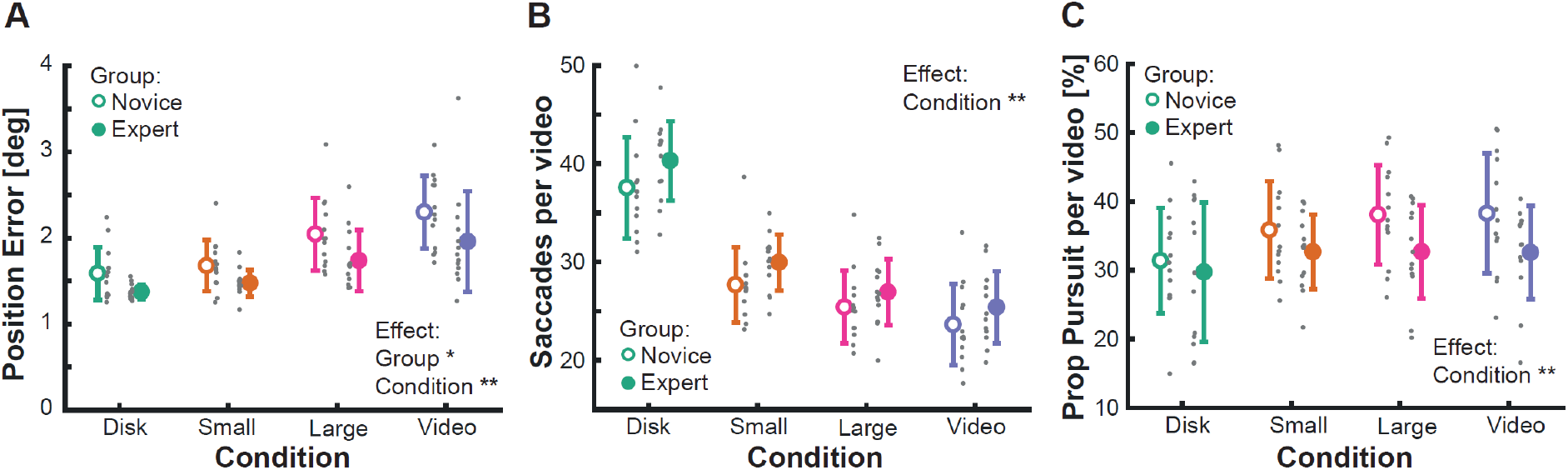
Overall eye movement behavior based on expertise and available peripheral information. **A** Average position error across the whole video is shown for the different conditions (different colors), separated for the level of expertise (open vs filled symbols). Circles show the average and standard deviation, gray circles show the individual data points. Note here that individual data points a slightly jittered horizontally for illustration purposes, but sometimes can still overlap. Since expertise was a between-subject factor, the data points cannot be matched to each other. **B** Average number of saccades per 10s video. Depiction as in A. **C** Proportion of time spend with pursuit eye movements per 10s video. Depiction as in A.

When investigating how many saccades and how much pursuit eye movements were used to track the puck, we observed an influence of the available peripheral information. The number of saccades (F(1.911,45.863) = 419.478, p < .001) as well as the proportion of pursuit (F(1.581, 37.954) = 8.153, p = .002) was significantly affected. There was no significant influence of expertise (Number of saccades: F(1,24) = 2.125, p = .158; Proportion of pursuit: F(1,24) = 2.287, p = .144) and no interaction (both p’s >.253). The number of saccades decreased with additional peripheral information (see Figure 2B, all comparisons across conditions: p < .001). This replicated our previous finding where we also observed roughly 30% less saccades from the disk to the video condition. At the same time, the proportion of the video spent with pursuit eye movements increased (see Figure 2C), as the disk condition led to significantly less pursuit than all other conditions (all p’s < .007), but no other comparison reached significance. This suggests that additional, more complex peripheral information leads to smoother tracking, with less saccadic eye movements and more time spend with pursuit.

#### Delay in tracking behavior

After characterizing general oculomotor behavior, we computed what we consider the most interesting metric: the estimated delay between eye and target movements. We estimated the delay via a cross-correlation (see Methods for details). Figure 3 shows the average cross-correlation across all observer for each of the respective delays, separately for novices (Figure 3A) and experts (Figure 3B). Overall, the pattern looks comparable: the disk condition led to the latest peak, whereas the peak shifted towards zero with an increasing amount of peripheral information. To quantify the effects, we estimated the peak of the cross-correlation for each observer and again computed an ANOVA. We observed a significant effect of peripheral information (F(2.184, 52.425) = 81.384, p <.001) and expertise (F(1,24) = 9.180, p = .006), but no significant interaction (F(2.184, 52.425) = 0.861, p = .437). Across all conditions, experts showed a significantly smaller delay when tracking the puck. For both groups, the delay reduced from around 160 ms in the disk condition to close to zero delay in the video condition (see Figure 3C). The differences between all levels of peripheral information were significant (all p’s < .006), suggesting an efficient use and integration of additional peripheral information and contextual cues that are available. Even only the little additional information that was present in the small condition, led to a significant improvement over the disk condition, suggesting that there were some cues around the puck (e.g., the skate movements of a player), that allowed better predictions. There was a lack of significant interaction between expertise and the available peripheral information, but experts reached the minimum delay novices showed when watching the full video already when only watching the video with the large cutout (see Figure 3C), and then kept improving further when having access to all available information.

**Figure 3.**
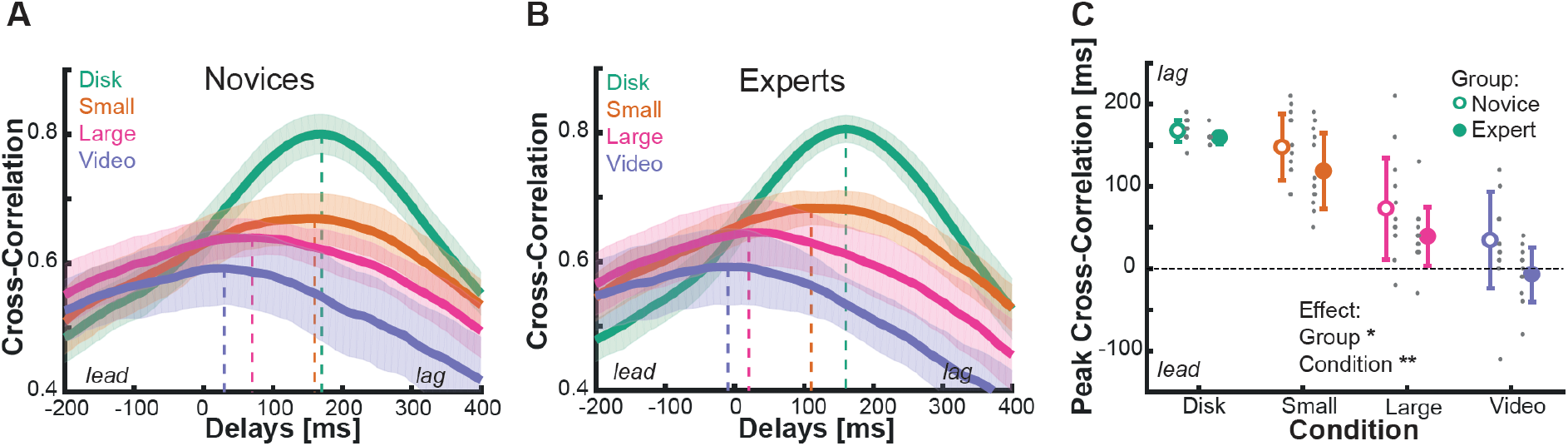
Estimated delay between eye and target movement. **A+B** Magnitude of the cross-correlation for different delays between eye and target movement averaged across all novices **(A)** or expert **(B)**. The vertical dashed lines indicate the peak of the cross-correlation, which serves as an estimate of the average delay between eye and target movement. The different colors indicate the different conditions. The shaded area shows the standard deviation across observers. **C** The peak of the cross-correlation for each observer averaged across the different conditions (color-coded) and level of expertise (open vs filled symbols). Circles show the mean and standard deviation, gray dots the individual data. Note here that individual data points a slightly jittered horizontally for illustration purposes, but sometimes can still overlap.

#### Delay for saccadic and pursuit eye movements

We observed a reduction of the average delay when observers had access to more information. Such an improvement could come from more predictive saccadic eye movements or could be achieved by more accurate pursuit. To address this point, in the next analysis we tried to estimate the contributions of saccadic and pursuit eye movements separately (see Methods for more details). For both, saccadic and pursuit eye movements (see Figure 4), we again found a significant influence of the available peripheral information (F(1.854, 44.502) = 167.527, p <.001 for saccades; F(2.713, 51.553) = 28.556, p < .001 for pursuit). For both eye movements we observed the same pattern: with increasing amount of peripheral information, the estimated delay decreased. However, although the metrics are not directly comparable (for saccades it represents the relative position of the saccade landing position to the target, for pursuit the estimate is based on cross-correlation over a longer time period), the overall difference in scale is interesting. It seems that even in the disk condition, saccades only land slightly behind the target, but land more and more ahead of the target with increasing information (see Figure 4A). In contrast, pursuit eye movements show the expected lag of around 170 ms in the disk condition, but then this delay decreases close to zero in the video condition (see Figure 4B). For the separate estimates of pursuit eye movements, we did not observe a significant effect of expertise (F(1,24) = 2.360, p = .141) nor an interaction (p = .357) with peripheral, although the pattern of results looks similar to the overall estimates (compare to Figure 3C). For saccadic delays we did observer an significant effect of expertise (F(1,24) = 5.378, p = .029) with experts showing lower delays across the conditions. In addition, there was also a significant interaction (p = .013), which was driven by comparable estimated delays for the disk condition, but then an increasingly larger difference between experts and novices the more peripheral information became available (see Figure 4A), suggesting that experts benefitted more from the additional peripheral information. The estimated delays for saccadic and pursuit eye movements were highly correlated across the different peripheral conditions (see Figure 4C, r(49) = .62, p <.001 for novices; r(46) = .67, p <.001 for experts), thus suggesting similar use of additional information for saccadic and pursuit control, although saccadic eye movements seem to be more strategically placed by expert observers.

**Figure 4.**
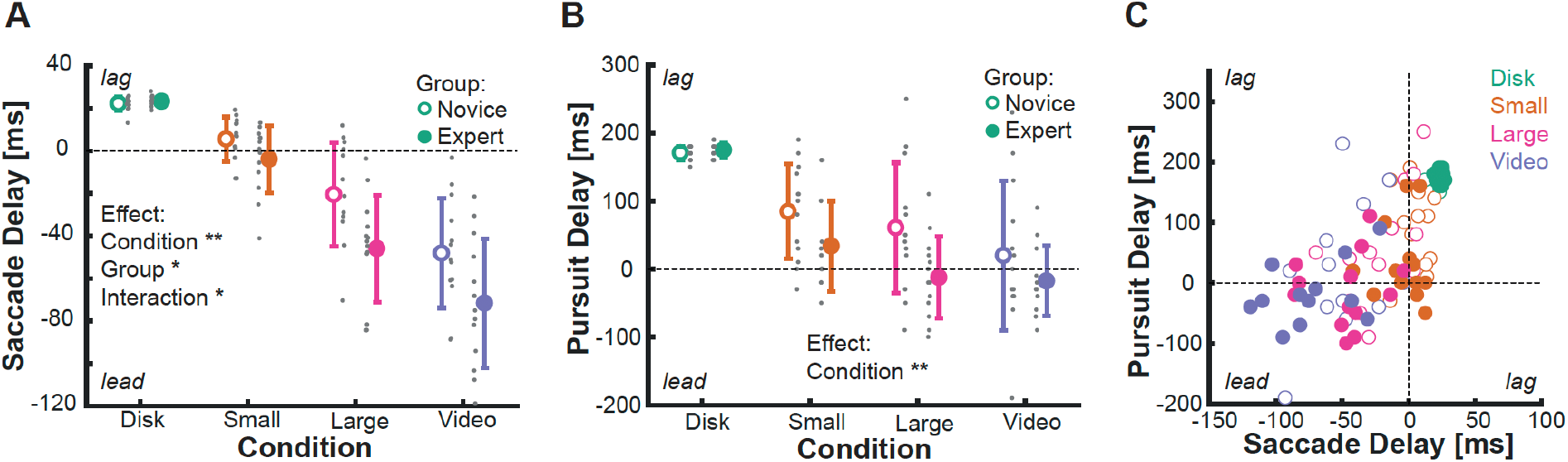
Separate estimate of delay for saccadic and pursuit eye movements. **A** The delay with respect to the target estimated for saccadic eye movements. Averages across subjects are shown for the different conditions (color-coded) and level of expertise (open vs filled symbols). Gray data points show individual data. Note here that individual data points a slightly jittered horizontally for illustration purposes, but sometimes can still overlap. Error bars depict the standard deviation **B** Same depiction as in A, but this time for the estimate of pursuit delay. **C** Estimated delay for saccadic and pursuit eye movements for each observer plotted against each other. Open symbols show data from novices, filled symbols data from experts. The color of the data points reflects the respective condition.

#### Passes as examples for prediction

To visualize predictive behavior better, we took a closer look at one specific situation in ice hockey games: passes between players. In a pass situation a successful prediction allows to already look at the receiving player before the puck arrives, which helps to analyze what is going to happen next. To quantify that, we computed when gaze arrived at the receiving player (see Figure 5A). We found again a significant impact of peripheral information (F(1.842, 44.204) = 102.481, p < .001) as well as a significant influence of expertise (F(1,24) = 5.807, p = .024), with no interaction (p = .726). The effects were again in the expected directions. Gaze arrived earlier at the targeted player for experts as well as with additional peripheral information (significant differences across all conditions with p<.001).

**Figure 5.**
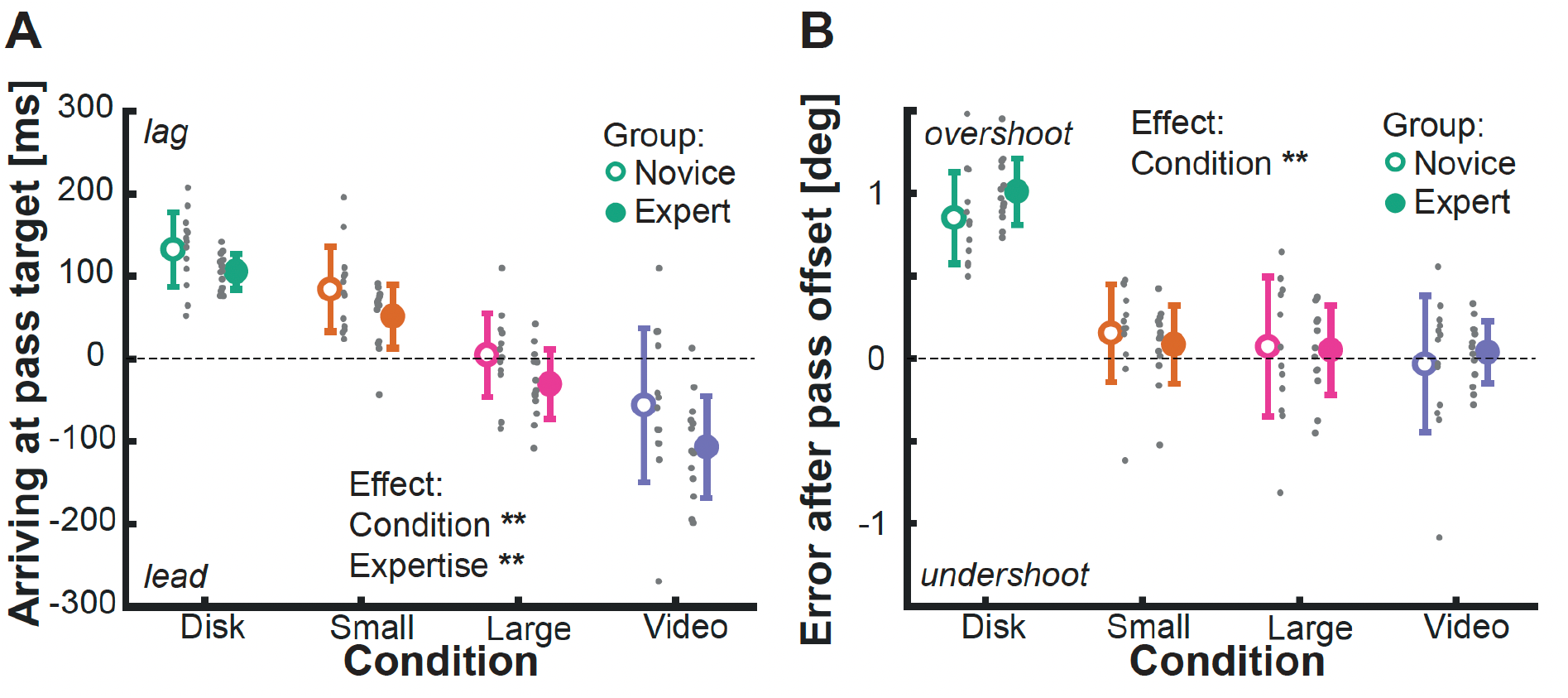
Eye movement behavior in pass situations. **A** The average time observers arrived at the pass target are shown for the different conditions (color-coded) and level of expertise (open vs filled symbols). Gray data points show individual data. Note here that individual data points a slightly jittered horizontally for illustration purposes, but sometimes can still overlap. Error bars depict the standard deviation. **B** Directional Error after pass offset. Negative values reflect an undershoot of the pass distance, positive values an overshoot. Depiction same as in A.

**Figure 6.**
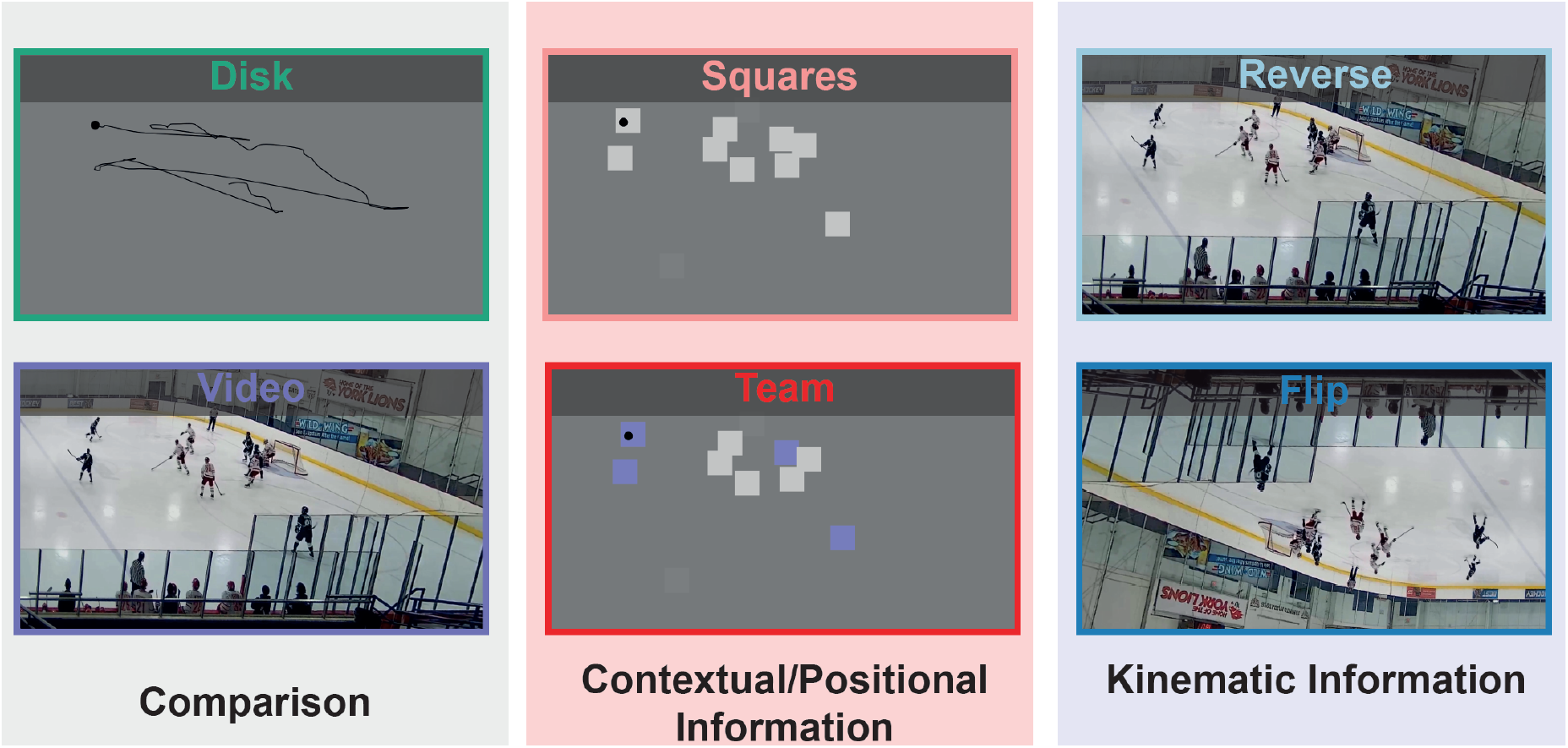
Conditions of Experiment 2. Depiction of the six different conditions. Disk and video conditions were identical to experiment 1 and our previous study. As an example, the whole trajectory of the puck is illustrated in the disk condition, please note that this was never visible to the observer. For the contextual conditions, the player positions were replaced by squares to provide positional information about the players. In the Team condition, the same information was presented, but additional information about the player identity was available. For the kinematic condition, the original video was shown, but this time either temporally reversed or flipped vertically. This manipulated the reliability of kinematic information about player movement, and especially in the reverse condition even changed the causal structure of movements. All six conditions were performed by each observer in random order.

These results suggest that observers were able to use the very little additional context in the small context condition to make some predictions about where the pass will go. However, this little context should only help, when the targeted player becomes visible. To check for that, we computed the average arrival time across observers for each of the labelled passed, based on the distance of the pass on the screen. We then computed linear regressions between the distance of the pass and the arrival time for each individual observer and condition. We found that the disk and video condition mainly differed in the intercept, but had on average very little slope. This is an interesting finding for the video conditions, since it suggests that even information in the far periphery can be integrated successfully for predictive eye movement. The two intermediate conditions clearly showed an influence of the pass distance. Both conditions showed positive slopes indicating that the larger the distance of the pass, the lesser the predictive advantage. This can be expected since for long pass distances, the targeted player simply only became visible quite late.

Observers did not only use the contextual cues available to arrive at the targeted player earlier, they also arrived there more accurately (see Figure 5B). When we computed the directional error 200 ms after pass offset (see Methods for details), the effect of expertise disappeared F(1,24) = 0.160, p = .693), but we still found a significant influence of peripheral information (F(2.754,66.096) = 95.395, p<.001). This suggests, that novices and experts were able to identify the targeted player at this point and fixated at them, but this was only possible for conditions with naturalistic information (disk vs other conditions p<.001). However, even the small context provided enough available information (all other comparisons p>.210) for an accurate fixation after pass offset.

## Experiment 2 - The role of contextual and kinematic cues

### Methods

Setup and Data Analysis are as in experiment 1

#### Observer

15 new observers (12 identified as female, 3 as male) took part in experiment 2. The age of the observers ranged between 19 and 54 years with a mean of 26.20 years. All participants were naïve to the purpose of the study and gave informed consent at the beginning of the experiment. The experiment was conducted in line with the Declaration of Helsinki and approved by the local ethics committee. Participants were paid 8 euro/hour for their participation.

#### Stimuli & Conditions

As in experiment 1 we again used 12 different 10s videos of ice hockey games across the different conditions. We kept the *disk condition* and the *video condition*, but since the focus of experiment 2 was the role of contextual and kinematic cues, four additional conditions were added.

To manipulate the contextual cues, we made use of additional labels of the position of players in the videos (Pidaparthy & Elder, 2019). In one condition (S*quares*) we replaced each player with a white square (Contrast = 0.6, Size = 1 deg), so that positional information about the rough position of the players was available, but no information about their team or their kinematic cues like body-movements. To increase the contextual cues, we also had a condition (*Teams*) with squares, but this time they were colored according to the team of their respective identity (Players with dark jerseys were shown in blue, the white team in white and referees in light gray).

To manipulate kinematic cues, we showed observers the original hockey videos, but one time flipped vertically (*Flip*), or in reverse (*Reverse*). Both conditions made it more difficult to process the kinematic cues of player movements, but especially the reverse condition was difficult, since it also changed the causal structure of puck movements, e.g., when a pass between players started, the player who seemed to start the pass did not perform a physically correct movement at this time.

Again, across all conditions observers were explicitly told that their task was to follow the target (disk in the disk or contextual conditions or puck in the video and kinematic conditions). Participants started each trial by pressing a space bar when looking at a central fixation cross, which was then followed by a video for 10s. At the end of the video, a small break occurred where the next video was prepared.

All observers completed all six conditions in random order. In each condition observers finished three blocks and within each block all 12 sequences were randomly presented. Each participant completed 216 trials (6 conditions x 3 blocks x 12 videos). One block took roughly 5-7 minutes and participants completed all 18 blocks in two 1-hour sessions, typically separated by multiple days. Across all participants this led to a total of 3240 trials (15 observers x 216 trials).

#### Exclusion criteria and statistical analysis

Exclusion criteria were identical to experiment: (1) if during the trial data was missing for more than 500 ms in a row and (2) the computed average error was larger than 10 deg. Based on these criteria 3141 of 3240 trials (92 %) were included in the analysis.

We computed the same metrics and averages as for experiment 1 and then statistical analysis was performed in JASP. Repeated-measure ANOVAs with the one factor (Disk, Squares, Team, Flip, Reverse, Video) were computed. If the sphericity assumption was violated, degrees of freedom were Greenhouse-Geisser corrected and corrected values are reported. To assess differences between the conditions, post-hoc t-tests were used. The p-value was adjusted according to Bonferroni-Holm to correct for multiple comparisons.

## Results

The goal of experiment 2 was to investigate the influence of contextual cues and kinematic information on predictive eye movements. We will again start reporting metrics related to overall eye movement behavior, and then look at the critical variable of the estimated delay between eye and target movement. To visualize the predictive behavior, we will end with looking at eye movement behavior in pass situations.

### Description of overall behavior

Overall, observers were more accurate in conditions when they could track a disk target (see Figure 7A) and had a gray background (all p’s <.001). The additional squares representing the player positions did not lead to a difference in the position error (all p’s >.165). While the kinematic conditions and the video condition overall had a higher position error, the unaltered video condition still had a significantly lower error than the conditions where we manipulated the video (both p’s <.001). A similar pattern of results was present for the number of saccades. The disk and contextual conditions had comparable number of saccades and were significantly different from the kinematic conditions and the original videos (see Figure 7B, all p’s <.001). However, the latter ones also had comparable numbers of saccades. For proportion of pursuit, there was substantial variability across observers with averages ranging from 10% to 60%. There was a significant main effect across all conditions (F(3.070, 42.980) = 3.255, p = .030), but only the comparison of the disk and the square condition with the video condition reached significance (both p’s < .026).There was a similar trend as in the first experiment: The video condition seems to led to a larger proportion of pursuit (M = 36.48 % +/-10.17) than the disk condition (M = 32.11 % +/-12.97).

**Figure 7.**
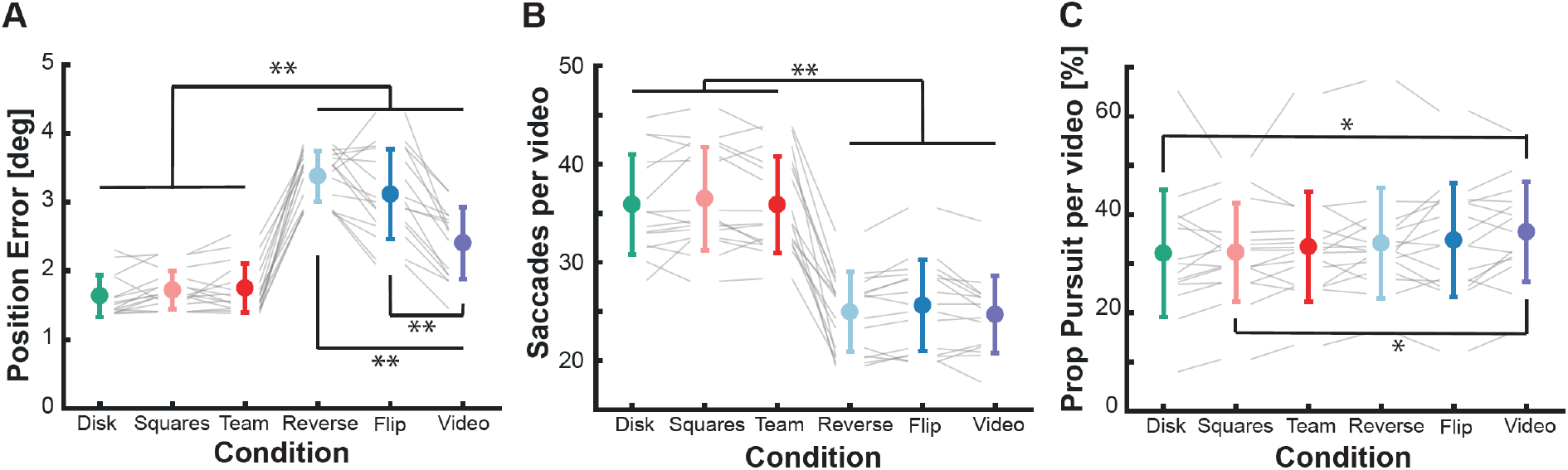
Overall eye movement behavior for different contextual and kinematic cues. **A** Average position error across observers shown across the different conditions (color-coded). Circles show the mean and standard deviation, gray lines show the individual subjects across all conditions. **B** Average number of saccades per video across the different conditions. Depiction otherwise as in A. **C** Average proportion of pursuit eye movements across the different conditions. Black lines indicate significant comparisons. ** p <.001, * p<.05.

### Delay in tracking behavior

After looking at some broader eye movement characteristics, we again look at the most interesting variable: the estimated tracking delay. As in experiment 1, we computed cross-correlations across varying delays across all conditions (see Figure 8A). To quantify the effect, we computed the peak of the cross-correlation for each observer and compared them across the conditions. There was a separation of the conditions into three different groups (see Figure 8B). The contextual conditions (squares and team) led to a comparable delay as the disk condition. The reverse condition was significantly worse than all other conditions with an average delay of 344 ms (SD = 58.895). The flip condition led to a similar delay as the regular video condition, which was significantly lower than the disk and contextual conditions. Therefore, the positional cues available in the contextual conditions did not seem to help to produce predictive eye movement behavior. If the causal structure of kinematic cues was impaired, the tracking delay became even higher than in the disk condition, where no cues were available. A vertical flip of the video, with intact causal structure but more difficult kinematic cues, still led to the significant prediction advantage we observed for the original videos.

**Figure 8.**
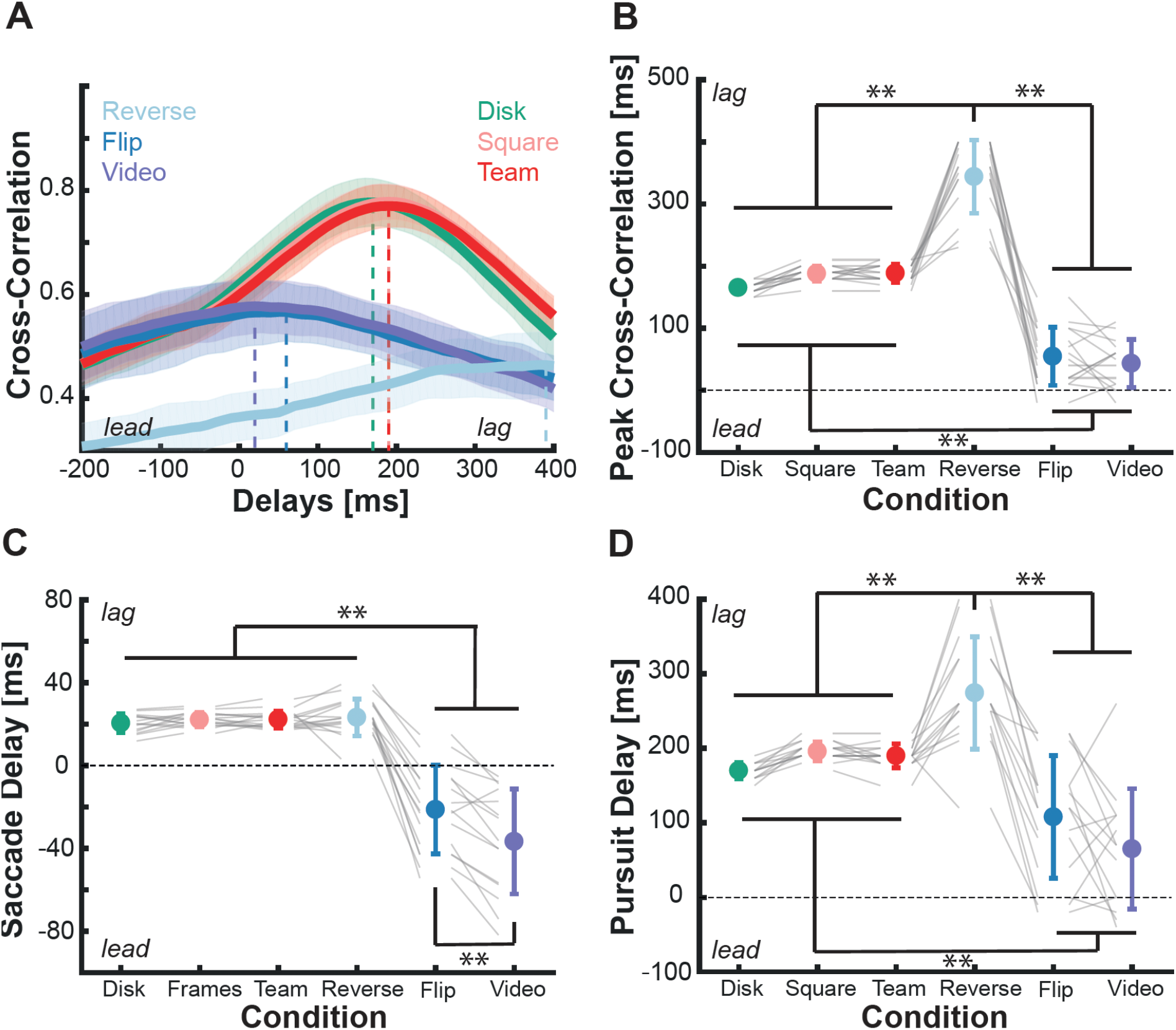
Tracking Delay for different contextual and kinematic cues. **A** Cross-correlation for different delays between eye and target movement averaged across observers for each condition (color-coded). The vertical dashed lines indicate the peak of the cross-correlation, which serves as an estimate of the average delay between eye and target movement. The shaded area shows the standard deviation across observers. **B** The peak of the cross-correlation across the whole video for each observer and each condition (color-coded). Circles and error bars show the mean and standard deviation. Gray lines depict the individual values of observes. **C+D** Estimates of the delay for either saccadic (**C**) or pursuit (**D**) eye movements. Depiction otherwise as in B. ** p<.01.

As in experiment 1, we also tried to estimate the delay separately for saccadic (see Figure 8C) and pursuit eye movements (see Figure 8D) to investigate their respective contributions. We found comparable patterns for both eye movements; the contextual conditions led to a similar lag as the disk condition. The flip condition showed a similar predictive advantage as the video condition (although it was slightly lower than the video condition for saccades). Interestingly, the reverse condition showed similar estimates of saccadic delay, suggesting that saccades in this condition show a similar reactive behavior as for the disk. However, for the pursuit estimates the reverse condition showed again the highest lag, suggesting that the impaired structure of kinematic cues also led to an impairment of pursuit performance. It might be that to gain a predictive benefit for pursuit eye movements, it is critical to be able to make longer predictions into the future, which is difficult for the reverse condition.

### Passes as examples for prediction

To look at the predictions in more detail, we again analyzed pass situations. Comparable to the results for saccadic delays, we saw that observers looked at the pass target roughly 100 ms after the puck arrived (see Figure A), suggesting a reactive gaze movement. However, for the flip and original video condition, we again observed the predictive advantage. Similar to experiment 1, we observed that when comparing the disk and video condition, observers did not only used the contextual cues to arrive earlier at the target, but also more accurately (see Figure 9B). The contextual conditions, led to significantly lower overshoot than the disk condition, suggesting that in pass situations observed made use of the available positional cues. The flip condition produced a similar error as the contextual conditions. The reverse condition led to no systematic over or undershoot, potentially suggesting that the motion cue of a pass and the pass situation in general helped observer to find the puck and look and fixate at the targeted player.

**Figure 9.**
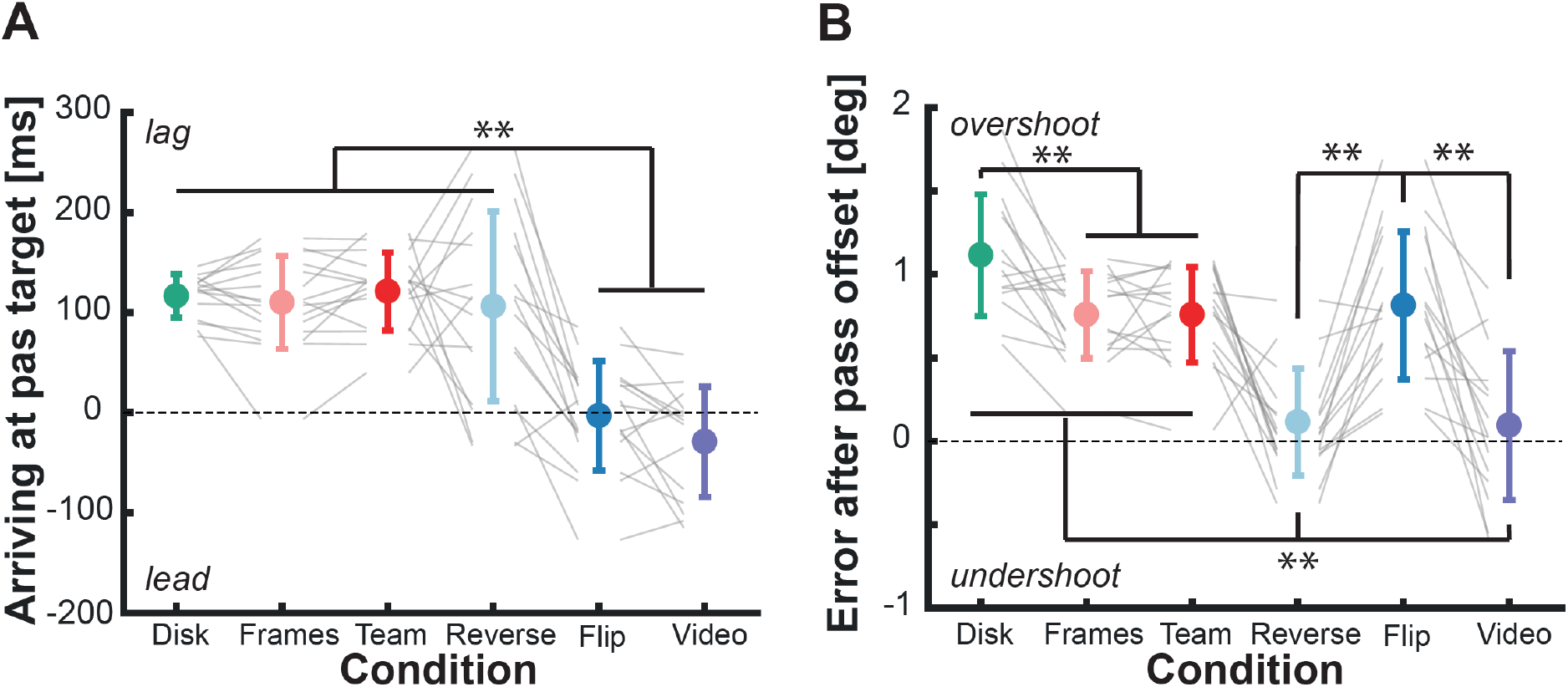
Eye movements in pass situations. **A** Average time observes arrived at the pass target across the different conditions. Circles and error bars show the mean and standard deviation, gray lines the individual values. **B** Directional error 200 ms after pass offset. Positive values reflect an overshoot, negative values an undershoot. Depiction same as in A. ** <.01.

## Discussion

A critical assumption in oculomotor research is that it is possible to generalize results obtained with simple, artificial stimuli to eye movement control in naturalistic, complex scenes. However, when we directly compared tracking performance for the same trajectories with simple stimuli and in complex naturalistic scenes (Goettker et al., 2020, 2021), we found that observers leverage additional information present in naturalistic scenes to achieve less tracking delay in their eye movements. But which kind of information is used by the oculomotor system to gain that predictive advantage? Here, we studied four potential influence factors: expertise, the amount of available peripheral information, and contextual and kinematic cues. Across most of our measurements, e.g., position error or predictive behavior in pass situations (see Figure 2A & Figure 5A), experts showed significantly better performance than novices. This suggests a role of experience with the complex stimuli, that allowed a better integration of the available information. In general, the more peripheral information became available, the better the predictions of all observers (see Figure 3), but experts seemed to be able to integrate them even more efficiently (see Figure 4A). When we manipulated the reliability of contextual and kinematic cues in the video, the most critical component seems to be access to intact kinematic cues from players. An artificial depiction of the player positions in the video did not lead to a predictive advantage (Figure 8) and only was of limited help in pass situations (Figure 9B). When the causal structure of kinematic cues was lost by playing the video in reverse, the tracking behavior was severely impaired, and the lag became even higher than without any contextual cues (Figure 8). Only the condition where kinematic cues were harder to process, but still intact due to vertical flip of the video led to similar predictive behavior that was observed when watching the original videos (see Figure 8). Together, these results demonstrate that when complex, naturalistic information is available, the oculomotor system is successfully integrating additional cues and is not only relying on low-level information about the target trajectory.

### The role of expertise

Across a broad range of research areas, the comparison of experts and novices has provided interesting insights into how experience changes behavior (Brams et al., 2019; Gegenfurtner et al., 2011; Memmert, 2009; Vickers, 2009). Especially in sports, there has been a focus on differences in eye movement behavior between experts and novices. For example, differences have been overserved in baseball (Land & McLeod, 2000), badminton (Abernethy & Russell, 1987), chess (Sheridan & Reingold, 2014, 2017), soccer (Casanova et al., 2009), table tennis (Ripoll et al., 1987) and many more situations (see Brams et al. (2019) for review). The general result seems to be that experts are better in identifying and predicting important information and therefore can allocate their gaze quicker to important locations (Brams et al., 2019). Although most of these findings were based on comparisons of active players, similar effects have also been seen when simply observing a game passively, which is more similar to the experiments we conducted (Smuc et al., 2010).

Our results match well into the current understanding of expertise effects. In our experiment, experts also showed lower tracking error and more predictive behavior than novices, which reveals a more efficient tracking strategy. For example, in pass situations experts are better in predicting where the pass will go. This allows them to already fixate the receiving player to analyze what is going to happen next. Similar observations have been made for other sports as well (Vansteenkiste et al., 2014; Williams et al., 1994). For saccadic eye movements, experts also benefitted more than novices from additional peripheral information (see Figure 4A), demonstrating a superior ability to make sense of and recognize potential peripheral cues, like the position of player or formations (Gorman et al., 2013), and use those for better predictions.

However, one important thing to note is that while the expert group outperformed the novice group, the novices also benefitted significantly from additional complex peripheral information and did quite well in the task. Across both experiments presented here, and our previous study (Goettker et al., 2021), when viewing the original clips of a high-speed sport as ice-hockey, novices were able to track the puck with on average close to zero delay. This clearly demonstrates that also novices can successfully integrate complex peripheral information to guide predictive eye movement behavior (Vig et al., 2011). The expertise effect, therefore does not lead to a fundamental difference in behavior, it is merely a domain-specific training of this predictive behavior that shows up with many years of experience in difficult situations (Tenenbaum et al., 2000). In naturalistic tasks and contexts, it seems to be a common feature of the oculomotor system to use knowledge about the world and task to perform predictive eye movements (Diaz et al., 2013; Goettker et al., 2021; Land & Furneaux, 1997; Land & Hayhoe, 2001; Sullivan et al., 2021).

### Integration of peripheral information

One interesting aspect of our experiment is that due to the task constraint of tracking the puck, experts might have even been at a disadvantage. A free-viewing study of soccer games, showed that experts in general seem to fixate more on the position away from the ball, like the position of other players, whereas novices fixated on the ball more frequently (Williams et al., 1994). Therefore, peripheral information became even more important in our task than it typically might be to predict what is going to happen next. Peripheral processing is a complex topic (Rosenholtz, 2016), but in general resolution drastically decays towards with the distance to the fovea (Grünert & Martin, 2020; Strasburger et al., 2011) with additional effects of clutter and crowding (Rosenholtz et al., 2007).

Our manipulations of the viewing window size revealed that cues for guiding predictive eye movements are available at multiple different distances. We observed a small predictive advantage when around 3 deg of the video around the puck position were available. A distance of 3 deg to the fovea is suggested to be critical area for the well-known research on the quiet-eye (Vickers, 2016). Quiet eye is defined as the ability to quickly bring and keep the gaze close to the target before an action towards it, which has been related to performance in different tasks. From a vision science perspective the importance of the central 3 deg seems rather arbitrary, since 3 deg away from the fovea resolution is already less than half of the foveal resolution and the eye is never really quiet (see (Spering & Schütz, 2016)). However, we think our results showed that giving observers access to 8 deg around the target allowed for a significantly better prediction. This adds direct evidence that information further out in the periphery is also used, and is important for guiding behavior. Since quiet eye is focused mostly on manual interception towards the fixated object, the use of such eccentric information to guide future eye movements to critical locations might not be directly comparable. However, making correct predictions about what is happening next is certainly highly important for our interactions with the environment. Additionally, we even observed an improvement in predictions when we showed the original video. Therefore, even information further out than 8 deg, seems to be useful. Thus, it seems that the oculomotor system is leveraging all available cues to improve tracking performance.

### The role of contextual and kinematic cues

Research on predictions in complex, naturalistic environments, such as sports, has mainly focused on two areas: kinematic cues and contextual factors (Loffing & Cañal-Bruland, 2017). Kinematic cues are related to information about the body movement of other players (which we manipulated with the flip and reverse condition), contextual cues are related to the knowledge about the position (which we manipulated in the squares and team condition) of an actor or their preferences.

Detecting and analyzing biological motion is a fundamental skill of human observers (Johansson, 1973; Neri et al., 1998). The high sensitivity of human observers to biological motion became visible when results showed that even only a few visible light-points attached to the joints of a walking human (so-called point-light displays), allow immediately to recognize a walking person. These limited displays even allow to make judgments about the sex (Kozlowski & Cutting, 1977) or emotional state (Atkinson et al., 2004) of the walker. The ability to use biological motion cues to make correct predictions about future behavior has been extensively studied in the sport context (Casanova et al., 2009; Loffing & Cañal-Bruland, 2017; Müller & Abernethy, 2012). For example, tennis players can use the movement of the opponent to make predictions about where they will hit the ball (Shim et al., 2005). In our study we decided to manipulate the quality of available kinematic information. Research with point light-walkers have shown that it is more difficult to recognize movements that are either played in reverse (Pavlova et al., 2002) or inverted (Pavlova & Sokolov, 2000). In the flip condition it was more difficult to analyze the biological motion cues, but the structure of the scene was otherwise intact. For the flip condition even naïve observers in experiment 2 could leverage the kinematic cues to gain a comparable predictive advantage as for the original video (see Figure 8). Interestingly, when we played the video in reverse, individual kinematic cues were more difficult to interpret, but also the causal structure of the movements was lost. For example, in a pass situation, the hitting movement of the stick did not start a pass, but actually was now only visible after the player received a pass with no consequences. This impairment of causality really seemed to confuse the observes reflected in only reactive saccades (Figure 8C & 9A), and overall, a much higher tracking lag than in the disk condition without any additional cues. Unexpected and surprising events are known to increase fixation durations (Vo & Henderson, 2009), which in dynamic scenes could be related to higher tracking delays. Together this demonstrates, that even naïve observers can successfully use kinematic cues to guide their eye movements. The underlying biological motion processing can deal with more difficult situations (e.g. an unnatural orientation), but the use of these cues breaks down when there are conflicts with our expectations of causal structure and our understanding of intuitive physics (McCloseky, 1983).

As mentioned above, next to kinematic cues, a second helpful source of information can be contextual cues (Loffing & Cañal-Bruland, 2017) or other situational information (Schläppi-Lienhard & Hossner, 2015). Previous research has demonstrated in squash that experts are able to predict above chance where an opponent will hit the ball, even before the opponent actually starts to move (Abernethy et al., 2001). Such predictions can be based on the position of the opponent (Loffing & Hagemann, 2014), domain and situational specific knowledge (Schläppi-Lienhard & Hossner, 2015), or information about the structure and organization of other players on the field (Casanova et al., 2009; Williams, 1993; Williams et al., 2006). We aimed at providing such contextual cues by showing information about the player position as simple squares. However, in both contextual conditions and whether the team identity of the players was visible or not, the contextual cues led to no predictive advantage in eye movement behavior. Across all metrics, performance was very similar to the disk condition, where no additional cues were present. Only in the error directly after a pass, there was a slight improvement (see Figure 9B). This could mean that observers were able to use the positional cues to fixate more accurately at the player position after the pass. Why the contextual cues did not lead to a predictive advantage in our experiment could be based on three reasons.

First, since the player positions were detected automatically (Pidaparthy & Elder, 2019) and were not manually labelled, they were not perfect. Especially when players were close together, occasionally one of the players could disappear. Additionally, since we made the 2D sizes of all squares constant, the 3D sizes of the players at different distances was not accurately represented. This might have decreased the reliability of the perceived player positions and led to a lower weighting of this potential cue. At least, they didn’t seem to serve as a distractor, since observers still achieved the same performance as in the disk only condition. Second, most of the studies reporting contextual effects investigated experts in the respective sport. Since in our second experiment, observers were again novices, it could have been that they simply were not able to make sense of certain player configurations and use those to their advantage. Third, as the result for the kinematic cues already shows, most of the predictive advantage seem to be based on the analysis of player movements to predict where they will go next. Also, the stick of the players, which was especially critical to predict the occurrence and direction of a pass was never visible. Together it seems that kinematic cues are just much more relevant for more precise predictive eye movements in our task, since they include the information about the future direction of a player movement, or the direction of a pass.

### Oculomotor control in naturalistic scenes

Across all conditions, the most striking differences in eye movement behavior was how tracking delay changed depending on expertise, available peripheral information and contextual and kinematic cues. However, besides the tracking delay there were substantial other differences. For example, the number of saccades decreased significantly when the puck was embedded in naturalistic context. In our previous study, we interpreted this effect as the need for less small corrective saccades due to better predictions (Goettker et al., 2021). However, based on these new results we must revise that statement, since also in the reverse condition (see Figure 7), where tracking delay was the highest across all tested conditions, the number of saccades was also significantly reduced. Therefore, the lower number of saccades is presumably more related to the complex visual input and in turn the less clearly defined position targets (Heinen et al., 2018). The reduction in number of saccades was often accompanied by more active pursuit eye movements (see Figure 2). Also, the delay estimates separately for saccadic and pursuit eye movements were correlated (see Figure 4C), which suggests a tight coupling between the two movements (Goettker & Gegenfurtner, 2021; Krauzlis, 2004; Orban De Xivry & Lefèvre, 2007). However, saccadic eye movements seem to be used in particular to bring the gaze to locations that are anticipated to become important in the near future (see Figure 4A & 8A).

All together our results indicate that eye movements leverage all available information around the target, experience and kinematic cues to reduce tracking delay. Instead of only using low-level sensory input to explain oculomotor control (Lisberger, 2010, 2015) or gaze selection (Itti et al., 1998), our results demonstrate the importance of higher-level information available in naturalistic contexts. We want to emphasize that while high-level information becomes more important in naturalistic and complex tasks, this does not render low-level information useless. A successful model needs to incorporate both, low and high-level factors (Torralba et al., 2006), and most predictions are based on a combination on both types of information (Badler & Heinen, 2006). For example, to make predictions about the future movement direction of a player, the correct analysis of biological motion is important, in the same way the extraction of the direction of a pass is critical to determine where it will go to anticipate the receiving player.

Our results provide empirical evidence for a suggestion made by Henderson (Henderson, 2017): In naturalistic scenes and tasks, gaze control is mainly based on predictions. While during some times the eyes are on a longer leash, and the deviation from the puck position is even slightly higher (see Figure 2 & 7; although note here that this error is also driven by predictive eye movements), the higher-level information is used to guide the gaze to crucial positions at the right time or even ahead of time (e.g., a player receiving a pass). This allows a better analysis of critical information to predict what is going to happen next. Such a strong influence from higher-level factors can be easily seen when looking at the selection of fixation points: While during free-viewing of a scene, predictions based on low-level image properties have been highly successful (Itti et al., 1998; Kümmerer et al., 2022), a simple task can render these predictions useless (Einhaeuser et al., 2008; Rothkopf et al., 2016; Tatler et al., 2011; Yarbus, 1967). Gaze control in natural tasks is more focused on fulfilling certain task demands, for example guiding an action or locating an object (Diaz et al., 2013; Hayhoe & Ballard, 2014; Land & Hayhoe, 2001; Sullivan et al., 2021), and therefore guidance by low-level factors becomes less important.

## Data Availability

Raw and aggregated data as well as stimulus information can be found at https://osf.io/kqup8/. Analysis is available at: https://github.com/AlexanderGoettker/CuesforNaturalisticVideos.

## Author contributions

Conceptualization A.G., N.B., L.L. & K.G.; methodology, A.G.; data collection, N.B. & L.L.; formal analysis, A.G.; writing—original draft, A.G.; writing—review & editing, A.G., N.B., L.L.. & K.G.; visualization, A.G.; funding acquisition, K.G.

## Acknowledgements

We want to thank Avi Aizenman for helpful comments on the manuscript. All authors were supported by the Deutsche Forschungsgemeinschaft (DFG; project number 222641018– SFB/TRR 135 Project A1).

